# Biophysical analysis of *Gaussia* Luciferase bioluminescence mechanisms using a non-oxidizable coelenterazine

**DOI:** 10.1101/2022.10.01.510427

**Authors:** Kyoko Takatsu, Naohiro Kobayashi, Nan Wu, Yves L. Janin, Toshio Yamazaki, Yutaka Kuroda

**Author notes:** **Correspondence:** YK and TY. Equal contribution (Kyoko Takatsu, Naohiro Kobayashi). **Data deposition:** The expression vector for GLuc-C9D (p21GLucTG) is deposited in Addgene (ID:124660). College of Food and Biological Engineering, Zhengzhou University of Light Industry, 136 Kexue Road, Zhengzhou 450002, P. R. China.

## Abstract

*Gaussia* luciferase (GLuc 18.2kDa; 168 residues) is a marine copepod luciferase that emits a bright blue light when oxidizing coelenterazine (CTZ). It is a helical protein where two homologous sequential repeats form two anti-parallel bundles, each made of four helices. We previously identified a hydrophobic cavity as a prime candidate for the catalytic site, but Gluc’s fast bioluminescence reaction hampered a detailed analysis. Here, we used azacoelenterazine, a non-oxidizable coelenterazine analog, as a probe to investigate its binding mode to GLuc. Interestingly, the biochemical studies of GLuc inhibition by azacoelenterazine also led us to find that salt, and monovalent anions, are required for GLuc’s bioluminescence, which seems reasonable for a sea-dwelling creature. The NMR-based investigation, using chemical shift perturbations monitored by ^15^NH-HSQC, suggested that CTZ binds to residue in or near the hydrophobic cavity. Of note is that these NMR data are in line with a recent structural prediction of GLuc, hypothesizing that large structural changes occur in regions remote from the hydrophobic cavity upon the addition of CTZ. Interestingly, these results point toward a unique mode of catalysis to achieve CTZ oxidative decarboxylation.

## Introduction

Bioluminescent organisms are found in more than half of all phyla in the animal kingdom (Shimomura and Yampolsky, 2016) and over 100,000 species in 13 phyla and 660 genera, from bacteria to bony fishes (Wu et al., 2003). Seventy-six percent of the 350,000 deep sea-dwelling organisms living are endowed with bioluminescence (Martini and Haddock, 2017). Bioluminescence is traditionally classified as fluorescence or chemoluminescence. In the latter class, the light signal is produced by a chemical reaction resulting from a luciferin – Luciferase pair. The term Luciferase denotes the enzyme catalyzing this reaction and luciferin describes small chemicals acting as their substrates (calcium-dependant Luciferases are also called photoproteins).

Thus, Luciferase is a generic term whose members are not necessarily evolutionary unrelated. Some luciferases require ATP or metal ions for activity, but not all. The Luciferase genes (or photoprotein genes) have been cloned for more than 60 species belonging to ten different groups of organisms. Similarly, luciferins do not necessarily have a common chemical structure (Shimomura and Yampolsky, 2016). Ten types of luciferins are known: Dinoflagellate luciferin (Nakamura et al., 1989), Bacterial luciferin (Strehler and Cormier, 1954), Cypridina luciferin (Kishi et al., 1966), Firefly luciferin (White et al., 1961, McElroy, 1947), Luminescent earthworm (Ohtsuka et al., 1976), shad shell latia (Shimomura and Johnson, 1968), Luminous mushroom (Purtov et al., 2015) and Coelenterazine (Inoue et al., 1975).

In the sea, a widely present class of luciferin are coelenterazine (CTZ) or some derivatives. It is indeed found in at least seven species of marine luminescent organisms (radiolaria, cnidaria, combs, copepods, head-foot, decapoda, and chaerognatha) (Widder, 2010). Moreover, if coelenterazine is readily oxidized by their respective Luciferase, there is no strong sequence similarity or even sequence motif common to these luciferases.

*Gaussia princeps* is a bioluminescent copepod which dwells in tropical or deep subtropical seas. Copepods are represented by more than 12,000 species and dominate the marine zooplankton fauna. Several bioluminescent copepods beside *Gaussia* have been reported: *Metridia longa* from the Arctic Ocean (Markova et al., 2004), *Metridia pacifica* from the Pacific Ocean (Takenaka et al., 2008), and over 20 bioluminescent copepods have been isolated around the Japanese coasts (Takenaka et al., 2012). The *Gaussia* luciferase (GLuc) gene was cloned in 2002 (Verhaegen and Christopoulos, 2002). Further studies pointed out common features for copepod luciferases such as a molecular weight of about 20 kDa, which is smaller than that of non-Caecilian luciferases, as well as an N-terminal secretory signal of about 20 amino acids. In addition, copepod luciferases contain two tandem repeats of about 60 amino acids, featuring highly conserved amino acids (especially Cys, Leu, Arg, and Pro. A consensus sequence for the copepod luciferase repeat is proposed as: C-x(3)-C-L-x(2)-L-x(4)-C-x(8)-P-x-R-C, where x stands for any amino acid (Takenaka et al., 2013). Like all copepods, luciferases GLuc use coelenterazine as their substrate (Campbell and Herring, 1990).

On the structural point of view, *Gaussia* luciferase (GLuc) is an all-alpha-helix protein made of nine helices as determined by heteronuclear multidimensional NMR (Wu et al., 2020). GLuc surface-accessible analysis indicated that 19 residues form a hydrophobic cavity, which was hypothesized to be involved in the bioluminescence activity. However, NMR experiments could not identify interactions between Gluc and coelenterazine by chemical shift perturbation as monitored by HSQC, possibly because of a fast turnover of the oxidation reaction.

In the present study, we analyzed the mechanism of GLuc-coelenterazine interaction using a non-oxidizable analog of coelenterazine which we named azacoenlenterazine (Aza-CTZ). We found that Aza-CTZ binds to or nearby the hydrophobic pocket of GLuc. Moreover, and less anticipated, we found that salt is essential for the bioluminescence of GLuc. Altough, retrospectively, this seems to be reasonable since luminous copepoda lives in halophilic environments, and thus, the natural reaction should occur in the presence of salt.

## Materials and methods

### Materials

Coelenterazine (CTZ) was purchased from Nanolight Technology. A stock solution of CTZ was prepared by dissolving CTZ in methanol at a concentration of 10 mM. The stock solution was diluted in the sample to the indicated final concentration for measurements. CTZ stock solutions were kept at -30 °C until use. Azacoelenterazine (Aza-CTZ) was synthesized in four steps from readily available 3-benzyl-5-(4-(benzyloxy)phenyl)-2-chloropyrazine (Coutant, E. et al., 2019) as previously described (Schenkmayerova, A. 2022) and this compound was dissolved in DMSO at a concentration of 42.4 mg / mL. We expressed the recombinant *Gaussia* luciferase (GLuc) in E-coli using a Solubility Enhancement Peptide tag (SEP-Tag), which was essential for producing natively folded GLuc as described previously (Rathnayaka et al., 2010, 2011, Wu et al., 2020)

### Luminescence assays

Light emission spectra were measured using an FP-8000 fluorescence spectrophotometer with an emission bandwidth of 5 nm and time response of 0.2 s (JASCO International Co., Ltd, Tokyo, Japan) at 15□. Measurements were initiated by the addition of CTZ to the samples.

For the salt effect experiments, 40 μl of CTZ solutions, prepared with various salts (NaCl, NaBr, NaI, NaF, KCl, CaCl_2_, MgCl_2_) to final concentrations of 1, 5, 10, 50, 100, 200, 500 and 1000 mM, were added to 10 ul of GLuc solution (final concentration of 0.5 uM GLuc, 25 mM MES pH 7.0). The mixture was thoroughly mixed by pipetting. The time between the substrate addition and measurement initiation was of 10 seconds.

### Inhibition assay

A mixture of CTZ and Aza-CTZ, where CTZ concentration was kept constant to 0.5 μM and Aza-CTZ concentration was varied from 0 M to 300 μM, was prepared. 40 μL of the CTZ+Aza-CTZ mixture containing a final concentration of 0.2 M NaBr and 0.3 M ascorbic acid was added to 10 μL of GLuc solution (final concentration of GLuc was 0.2 μM in 25 mM MOPS buffer, pH 7.9). The sample was mixed by five-time pipetting, and the spectrum was measured. The average maximum intensity measured when DMSO alone was added was 1400 (without Aza-CTZ).

### NMR analysis

NMR experiments were conducted using 0.1 mM ^15^N labeled GLuc protein dissolved in 50 mM MES buffer pH 6.0 and 2 mM NaN_3_, at 293 K with 8% (v/v) D_2_O in a 5 mm Shigemi microtube (Shigemi co., Ltd, Tokyo, Japan). NMR spectra were acquired on a Bruker Advance-III 600 MHz spectrometer equipped with a 5 mm CPTXI cryoprobe. Two-dimensional ^1^H-^15^N heteronuclear single quantum coherence (HSQC) experiments were performed for chemical shift displacement analysis upon the addition of Aza-CTZ.

## Results and Discussion

### The effects of ions type on GLuc luminescence

The recombinant GLuc dissolved in 50 mM Tris-HCl showed a strong bioluminescent activity upon the addition of CTZ, in line with our previous observations. However, under the same condition, but in 50 mM MES instead of Tris-HCl, Gluc was inactive. The addition of NaCl restored Gluc bioluminescence activity in MES buffer, prompting us to carefully investigate the effects of salts on GLuc’s activity. We thus measured the luminescence intensity catalyzed by GLuc using five monovalent anions, four halide ions (F^-^, Cl^-^, Br^-^, I^-^) and a nitrate ion (NO_3_^-^) (Fig. 1). The Cl^-^ and Br^-^ ions increased GLuc’s luminescence, and NO^3-^ did so to a lesser extent. The luciferase activity was highest between 50 and 150 mM, depending on the ion type. The addition of Cl^-^ and Br^-^ induced the strongest light emission, while I^-^ showed a weak emission, and F^-^ did not induce luminescence at all. These results are in line with a previous report (Inouye and Sahara, 2008a and 2008b). In the case of chloride ions, the use of NaCl, KCl, MgCl_2_ or CaCl_2_ solutions led to similar results.

**Fig. 1.**
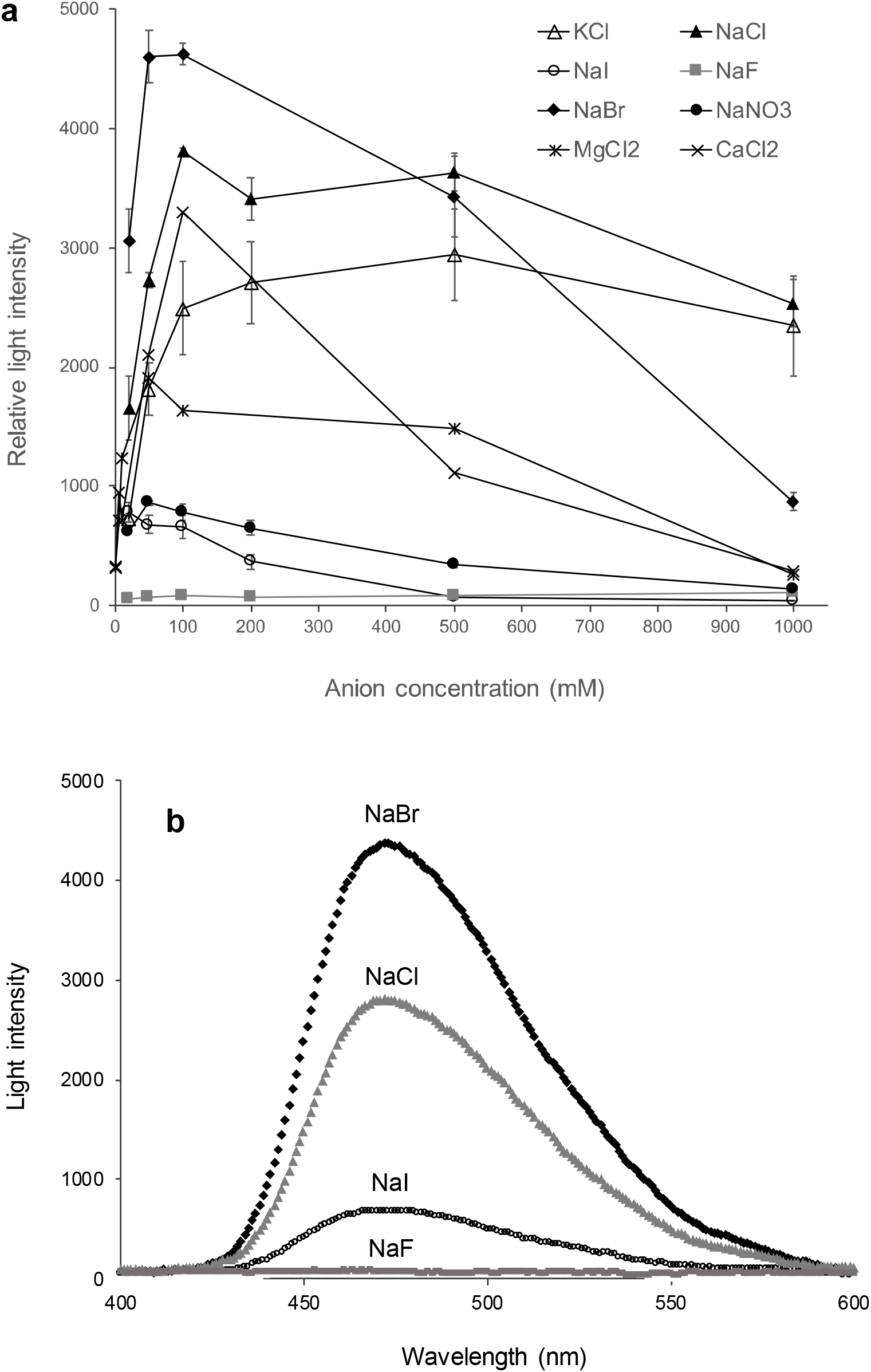
Effect of salt on the luminescence *Gaussia* luciferase. **(a)** In the luminescence assays, 5 μM of CTZ containing a salt to be tested was luminesced in the presence of 0.5 μM of GLuc in 50 mM MOPS buffer, pH 7.9, at 15 □. Salt concentration ranged from 20 to 1000 mM. Error bars indicate the standard deviation. (b) Effect of halogen ions on the luminescence spectra. The solution conditions are the same as in (a) but with a fixed salt concentration of 50 mM.

Dissolved monovalent halides content (F. Culkin, 1965) in the sea is high for chloride (0.546 mol/kg) but much smaller for bromine (0.844 mmol/kg) or fluorine (0.068 mmol/kg) and even less for iodine. Interestingly, only chlorine and bromine were found essential for this bioluminescence reaction. We hypothesized two reasons for the remarkable lack of activation by fluorine or the modest effect of iodine ions. First, if the anion is involved in a specific interaction such as a salt bridge, the anion’s size should be an issue. F^-^ is the smallest monovalent halogen wheras iodine has the biggest (anions’ thermochemical radius for Cl^-^: 1,81 Å, Br^-^: 1,96 Å, and I^-^: 2.20 Å and F^-^: 1.33 Å (Simoes et al., 2017)). A second hypothesis is that fluorine ions, which have the highest electronegativity, might repel each other and hamper their close contact in a cavity.

### Inhibition assay

We performed inhibition assays using Aza-CTZ (Fig. 2) and CTZ. To this end, we first assessed the solubility of Aza-CTZ with water, methanol and other alcoholic solvents as well as dimethyl sulphoxide (DMSO) to use it as a stock solution. Aza-CTZ turned out to be only suitably soluble in DMSO. DMSO is also known as a solvent used to trigger the chemiluminescence of CTZ (Teranishi and Goto, 1990), but as measured for Aza-CTZ, the baseline was low and indicated a maximum intensity of 1400. Indeed, Aza-CTZ is stable in DMSO since the carbon oxidized in CTZ is replaced by nitrogen. We then faced two problems with a stock solution of Aza-CTZ in DMSO. Firstly, the solution of Aza-CTZ did not completely dissolve in an aqueous solution above 150 μM. Accordingly, the final concentration of Aza-CTZ used in our experiements had to be below 150 μM. For this reason, the CTZ concentration also had to be lowered which caused a reduction of the luminescence intensity.

**Fig. 2.**
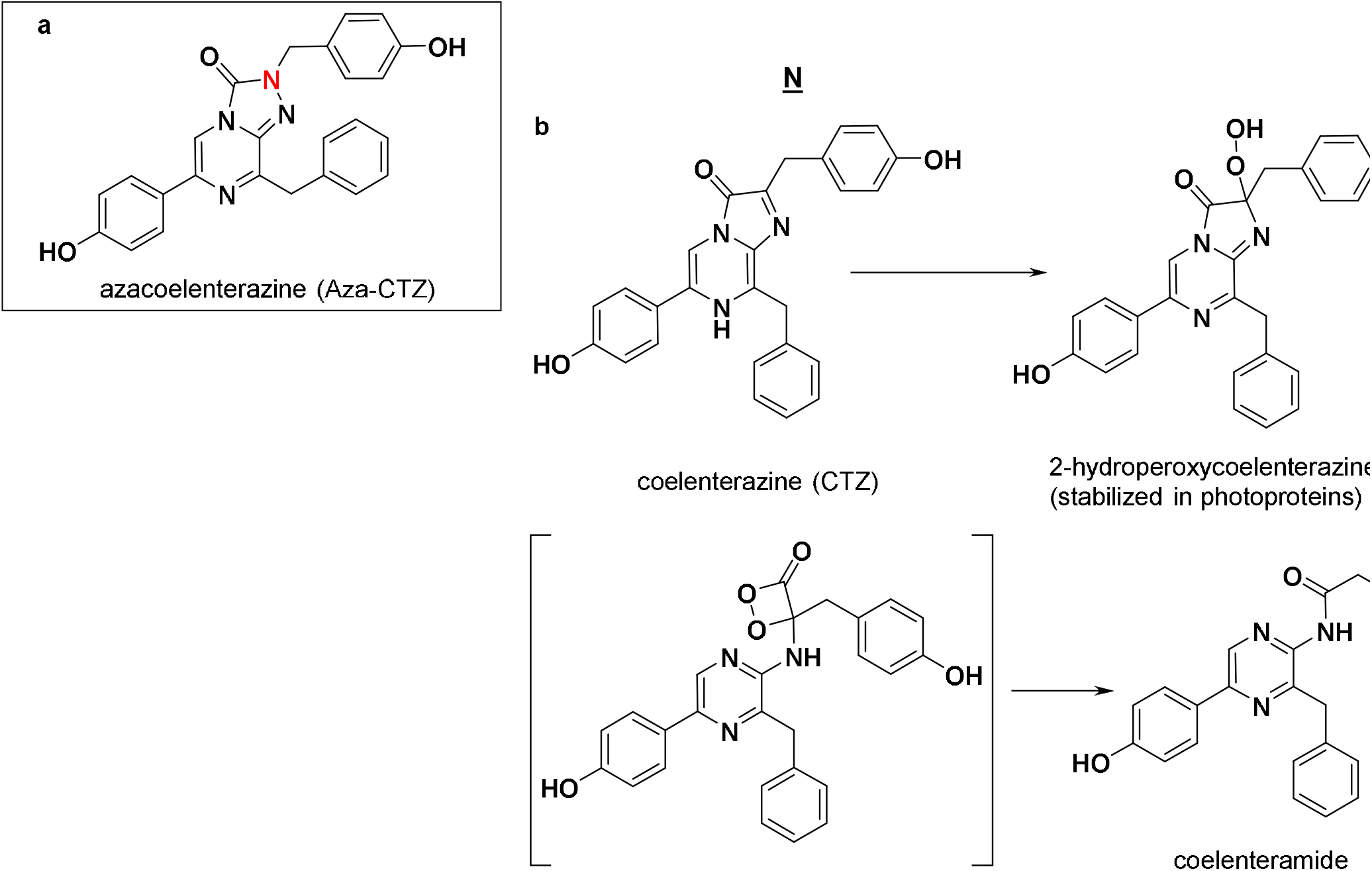
Chemical structure of Aza-CTZ and oxidative decarboxylation reaction of CTZ. (a): structure of Aza-CTZ showing the carbon replaced by a nitrogen atom by an underlined <u>N</u>. (b) two stages of the oxidative decarboxylation process of CTZ into coelenteramide leading to the production of a photon as well as CO_2_.

We then optimized the reaction pH using four buffers at different pHs: MES buffer pH 6.0, MES buffer 7.0, MOPS buffer 7.9, and Tricin buffer pH 8.8. NaOH was used to adjust the pH of buffers. We found that the luminescence intensity was strongly pH-dependent, but the wavelength of the intensity maximum remained unchanged. Eventually, MOPS buffer at pH 7.9, was selected as it led to the strongest bioluminescence. The second issue was that the addition of DMSO suppressed the bioluminescence intensity in our experiments. We found that the final DMSO concentration had to be under 1% to avoid this. In fact a bioluminescence reduction by DMSO has been previously observed with the CTZ-using *Renilla* luciferase, (Mishra et al., 2017). In addition, the reaction solution had to be mixed for precisely 10 seconds to increase the measurement accuracy. Thus, we eventually used ascorbic acid (vitamin C) as a reductant to slow down any oxydation reaction and improve the measurement accuracy. Thus, the inhibition assays were performed in 50 mM MOPS buffer, pH 7.9, 0.2M NaBr, containing 0.3 M vitamin C. Under these conditions, Aza-CTZ inhibited the bioluminescence intensity dose-dependently. A plot of the light intensity *vs*. Aza-CTZ concentration showed a sigmoidal curve (Fig. 3) with a 50% inhibition concentration (IC_50_) between 10 and 100 μM, equivalent to a CTZ/Aza-CTZ proportion of 1/20-200. Unfortunately, further refinement of this IC_50_ value could not be achieved because of the limitations mentioned above.

**Fig. 3.**
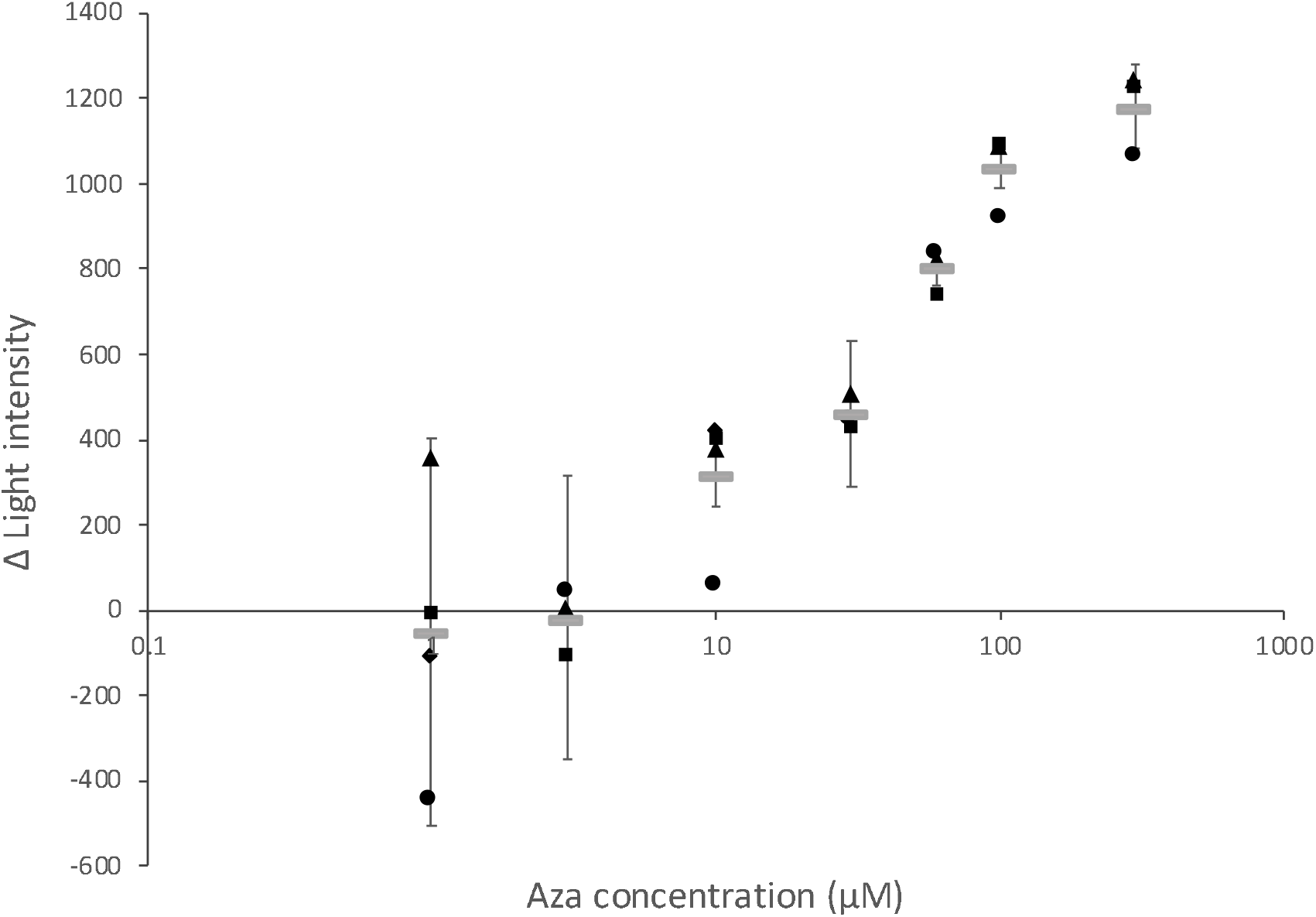
Inhibition assay. Relationship between the concentration of Aza-CTZ and the maximum intensity of luminescence. For inhibition assay, 40 μL of CTZ mixture containing 300 μM ∼ 1 μM or 0 M of Aza-CTZ was added to 10 μL of GLuc solution and pipetted five times. Then the spectrum was measured. The final concentration of CTZ and GLuc was 0.5 μM and 0.2 μM, respectively.

### NMR analysis

GLuc is an all-alpha-helix protein made of nine helices (Wu N., 2020). GLuc surface-accessible analysis indicated that 19 residues formed a hydrophobic cavity that might be involved in bioluminescence. The cavity is made by 19 residues around the central α1 (N10, V12, A13, V14, S16, N17, F18), α4 (L60, S61, I63, K64, C65), the functionally important but highly flexibke loop (R76, C77, H78, T79), and α7 (F113, I114, V117).

We did not observe chemical shift perturbation in the HSQC spectrum upon the addition of CTZ to the NMR sample, but a bright light appeared, disappearing within a few tens of seconds, which is too fast for initiating NMR measurements (unpublished observations). We speculate that the lack of chemical shift perturbation is due to the fast turnover of the reaction and that CTZ quickly dissociates from GLuc upon its decarboxylative oxidization.

Accordingly, we expected that Aza-CTZ, which GLuc cannot oxidize, might “freeze” the reaction in the bound state, enabling us to see chemical shift perturbation and consequently deduce a putative CTZ’s binding site. Chemical shift perturbation was observed for A13, N17, A19, T20, N46, A47, R48, F72, I73, R76, T79, V134, R147, F151, and A152 (Fig. 4 and 5). The residues in the central α1 and the functionally important loop are common with the cavity residues listed above. The other residues are located in the C-terminal loop (none in α7), which is not close to the cavity in the NMR structure, but might be folded in an α-helix replacing the N terminal helix according to a predicted structure (Huang YJ, et al. 2021).

**Fig. 4.**
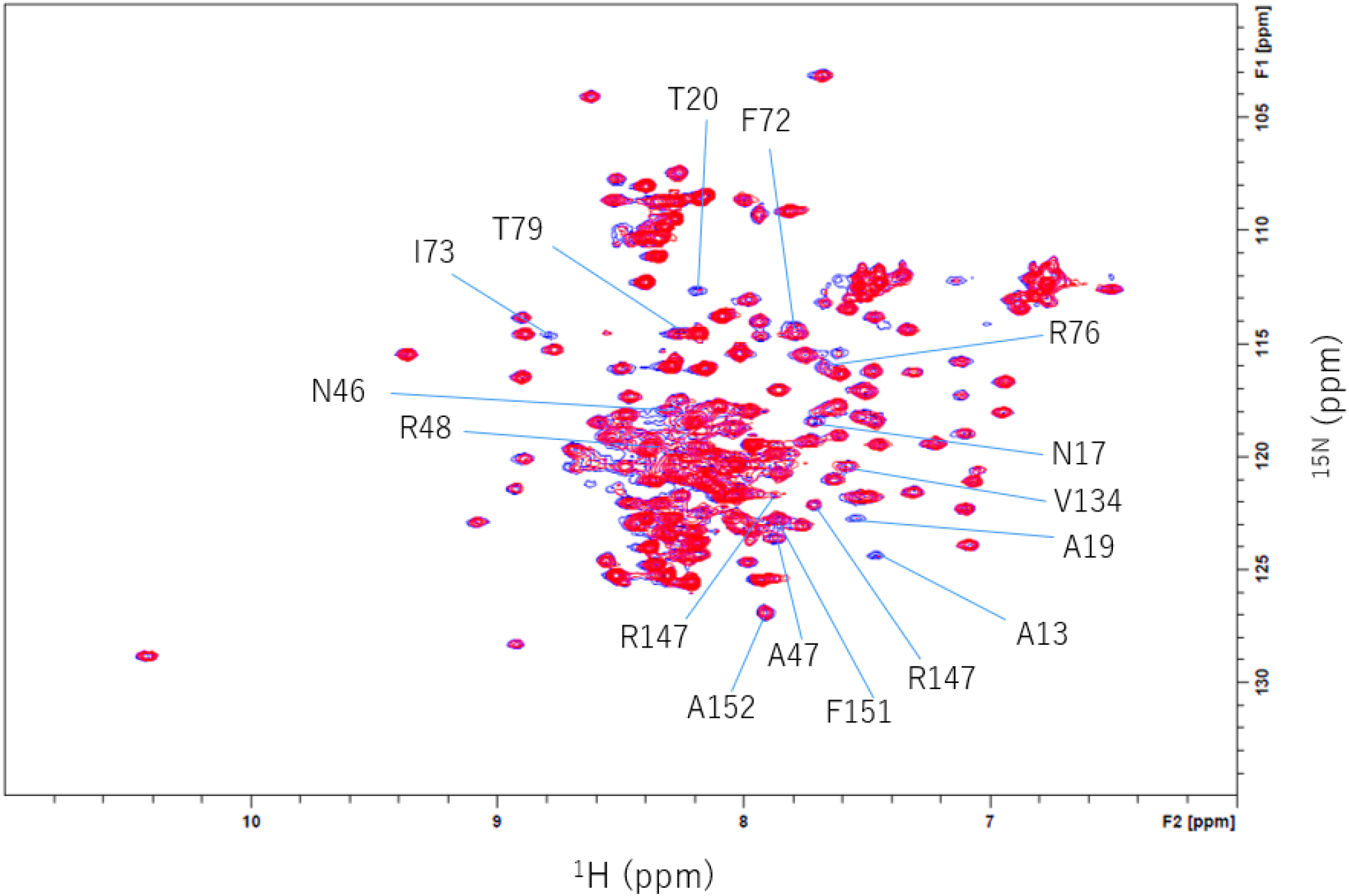
2D ^15^N-^1^H HSQC of GLuc in the presence and absence of Aza-CTZ. The spectra were measured **at 298K** with 0.2 mM aza-CTZ (1% DMSO) in red and without Aza-CTZ in blue. The ^1^H-^15^N peak positions were confirmed by 3D HNCO spectrum as the sequential ^1^H-^15^N-^13^CO (*i*-1) signals tend to be less sensitive for the little difference of experimental conditions. Upon addition of aza-CTZ, several peaks in the 2D ^15^N-^1^H HSQC spectrum were reduced, but no significant change in the chemical shift position was observed.

**Fig. 5.**
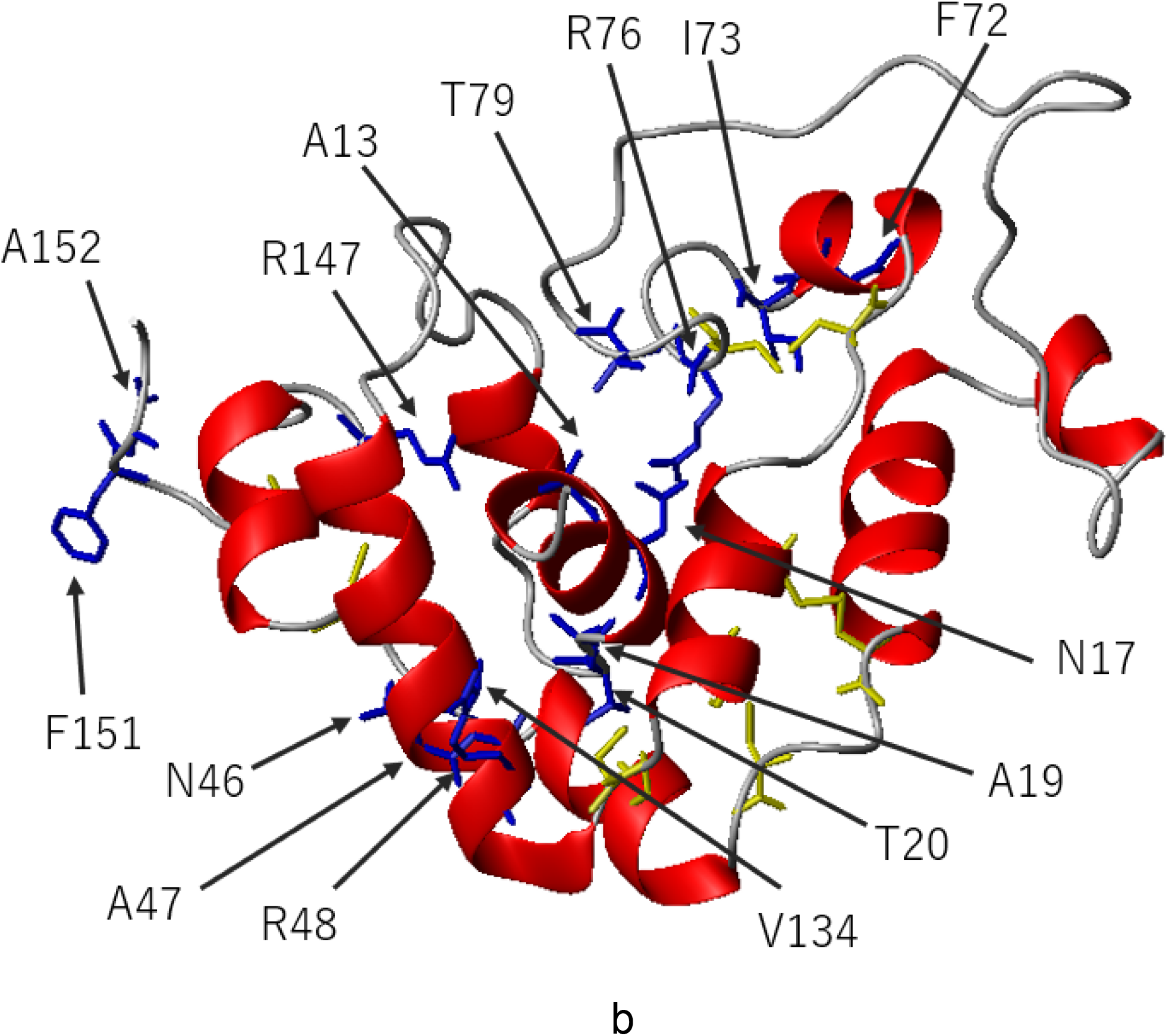
Ribbon model of GLuc. Ribbon model of GLuc showing the side-chains of the residues with significantly reduced ^15^N-^1^H HSQC peak intensities upon addition of Aza-CTZ (blue). SS bonds are shown in yellow. For convenience, the N-terminal residues 1-10 and C-terminal 150-168 regions are not shown.

Gluc contains several highly flexible or disordered regions [Residues P74 to A100 (including the functionally important loop) and P145 to the C-terminus]. We thus anticipated that the binding of CTZ to GLuc occur according to a fit-and-lock model possibly involving residues that are sequentially apart from the cavity rather than a key-and-lock model where CTZ would fit into the hydrophobic pocket. A fit and lock involving e a large structural change would be more in line with the observed substrate cooperativity (Tzertzinis G 2012) than a key and lock model where a single substrate fits into a preformed cavity. One such fator is the the alpha-fold predicted conformational change that involves unfolding the C terminal helix and its replacement by an N terminal helix (Huang YJ, et al., 2021)

Previous mutational analysis indicate that F72, I73, R76, H78, Y80 (Kim et al., 2011) and F113, I114, W143, L144, F151 (Wu et al., 2016) are activity-related residues. F72, I73, R76 and F151 were identified by our present chemical shift perturbation NMR experiment, strongly suggesting that these four residues are directely involved in the binding of CTZ (We suppose that the binding of mode of Aza-CTZ is identical to that of CTZ). No Chemical shift perturbation was found for H78, Y80, F113 and I114, but these four residues are located on or near the hydrophobic cavity identified in the NMR structure. We speculate that these residues contribute to the formation of the hydrophobic cavity without directly participating to the binding of CTZ. On the other hand., the effect of W143 and L144 (Wu et al., 2016), which are located in the unstructured C-terminus, is difficult to rationalize from the NMR sturcure. A structural rationale for their involvement in the bioluminecsce might be provided by the predicted structure (Huang YJ et al., 2013), where W143 and L144 would migrate closer to the hydrophobic cavity.

## Conclusion

We have analyzed the bioluminescence activity of *Gaussia* luciferase (GLuc 18.2kDa; 168 residues) using a recombinant GLuc expressed in *E* coli and refolded using a SEP tag as well as Aza-CTZ, a non-oxidizable coelenterazine analog. Firstly we found that salt is really necessary for GLuc’s bioluminescence. The optimum salts were Na and Br, at approximately 100 mM. Competition inhibition assay indicated that Aza-CTZ’s IC_50_ was between 10-100 uM. Secondly, we identified 15 residues whose peak intensity decreased upon adding Aza-CTZ suggesting that CTZ binds around cavity and the flexible functionally important loop. In addition, we speculated that Aza-CTZ binding causes a structural change, including the above-discussed one in the C-terminal region. Further structural analyses are needed to confirm whether such daring structural change occurs upon binding of the salt and/or CTZ.

## Note

The SCP-tag sequences are covered by a Japanese patent 5273438 and an international (PCT) patent application PCT/JP2018/029395.

## Authors’ Contributions

KT, YK, TY, and YLJ designed the project. KT, YK, and TY wrote the manuscript, YLJ and NW provided the materials. KT, TY, and NK performed the experiments and analyzed and compiled the data. All authors read and approved the manuscript.

## Data Sharing

All data are given in the manuscript and the supplementary data. The GLuc plasmid is deposited in ADDgene (ID: 124660).

