## supplemental for "Biophysical analysis of *Gaussia* Luciferase bioluminescence mechanisms using a non-oxidizable coelenterazine"

**Supplemental materials (2 figures)**

|  | pH 6.0 | pH 7.0 | pH 7.9 | pH 8.8 |
| --- | --- | --- | --- | --- |
| 400 | 27.9351 | 16.2543 | 46.0162 | 36.401 |
| 401 | 27.4893 | 16.3069 | 41.5245 | 37.9404 |
| 402 | 29.8041 | 10.7319 | 46.6561 | 35.1003 |
| 403 | 33.6435 | 15.8712 | 48.102 | 41.1307 |
| 404 | 30.139 | 9.84035 | 44.9905 | 39.3915 |
| 405 | 32.4584 | 10.9278 | 46.9837 | 46.0542 |
| 406 | 28.8344 | 7.36217 | 40.0981 | 45.5802 |
| 407 | 40.4067 | 14.7582 | 42.2345 | 43.3128 |
| 408 | 38.3267 | 15.9168 | 44.2203 | 43.7004 |
| 409 | 41.3304 | 18.0298 | 46.2686 | 42.1146 |
| 410 | 45.1885 | 22.0858 | 46.3487 | 39.474 |
| 411 | 38.419 | 15.7209 | 37.8492 | 43.2722 |
| 412 | 38.8265 | 17.6921 | 45.338 | 40.1852 |
| 413 | 40.3838 | 18.4243 | 47.9391 | 46.2835 |
| 414 | 40.0803 | 14.8495 | 45.2711 | 47.1838 |
| 415 | 43.171 | 22.5263 | 48.4864 | 49.6693 |
| 416 | 45.4572 | 26.8976 | 53.4137 | 57.1645 |
| 417 | 41.3775 | 26.2041 | 58.9706 | 54.6693 |
| 418 | 36.7561 | 21.2656 | 53.406 | 49.6864 |
| 419 | 39.3299 | 29.43 | 50.2306 | 53.286 |
| 420 | 42.9316 | 27.599 | 49.5111 | 53.508 |
| 421 | 43.0713 | 39.3316 | 54.2403 | 57.4477 |
| 422 | 36.2544 | 31.9495 | 52.4925 | 46.7962 |
| 423 | 42.4503 | 37.2347 | 58.4834 | 55.0254 |
| 424 | 38.4268 | 34.3271 | 57.9525 | 57.8643 |
| 425 | 47.6727 | 39.566 | 63.4936 | 65.3578 |
| 426 | 48.4644 | 37.9008 | 64.6222 | 64.0071 |
| 427 | 47.5925 | 37.7411 | 69.3196 | 69.7143 |
| 428 | 50.8441 | 41.6166 | 78.7183 | 76.5384 |
| 429 | 48.2259 | 48.8699 | 86.0638 | 76.1557 |
| 430 | 52.5624 | 59.0982 | 91.8186 | 80.8428 |
| 431 | 51.4028 | 64.6169 | 101.974 | 82.0338 |
| 432 | 52.4944 | 66.8409 | 103.586 | 85.503 |
| 433 | 60.9163 | 73.365 | 114.636 | 95.0373 |
| 434 | 62.0919 | 77.8951 | 130.148 | 109.233 |
| 435 | 67.0562 | 88.2365 | 146.237 | 123.058 |
| 436 | 72.5523 | 98.4966 | 162.167 | 137.744 |
| 437 | 82.9846 | 116.595 | 183.613 | 156.159 |
| 438 | 95.8574 | 132.736 | 204.668 | 176.502 |
| 439 | 105.602 | 161.353 | 233.663 | 195.431 |
| 440 | 120.729 | 187.942 | 269.919 | 210.345 |
| 441 | 123.379 | 205.596 | 296.358 | 232.986 |
| 442 | 124.312 | 226.098 | 328.837 | 256.621 |
| 443 | 135.761 | 259.767 | 367.785 | 287.641 |
| 444 | 151.775 | 291.887 | 410.419 | 321.006 |
| 445 | 177.592 | 323.203 | 449.039 | 356.153 |
| 446 | 208.784 | 364.437 | 499.061 | 393.454 |
| 447 | 231.193 | 404.004 | 553.58 | 434.609 |
| 448 | 261.794 | 447.317 | 604.677 | 472.538 |
| 449 | 279.348 | 482.811 | 662.642 | 507.225 |
| 450 | 299.019 | 529.088 | 712.762 | 547.867 |
| 451 | 334.665 | 587.771 | 783.875 | 601.833 |
| 452 | 361.648 | 637.165 | 856.416 | 646.459 |
| 453 | 382.078 | 678.698 | 920.179 | 684.539 |
| 454 | 407.757 | 724.993 | 983.64 | 727.699 |
| 455 | 435.593 | 775.908 | 1047.13 | 772.069 |
| 456 | 461.981 | 817.529 | 1107.51 | 814.639 |
| 457 | 496.724 | 879.702 | 1174.73 | 872.357 |
| 458 | 526.308 | 933.064 | 1245.32 | 908.238 |
| 459 | 554.387 | 981.87 | 1304.87 | 957.346 |
| 460 | 585.654 | 1031.06 | 1361.69 | 1000.37 |
| 461 | 604.579 | 1072.46 | 1414.05 | 1030.63 |
| 462 | 622.186 | 1116.55 | 1473.9 | 1066.18 |
| 463 | 630.672 | 1148.41 | 1514.58 | 1095.27 |
| 464 | 644.641 | 1184.24 | 1559.29 | 1118.49 |
| 465 | 666.605 | 1222.82 | 1599.22 | 1148.06 |
| 466 | 678.548 | 1245.71 | 1627 | 1170.39 |
| 467 | 692.692 | 1272.61 | 1650.85 | 1184.78 |
| 468 | 694.326 | 1287.91 | 1655.08 | 1196.19 |
| 469 | 704.738 | 1308.82 | 1667.85 | 1213.78 |
| 470 | 712.907 | 1328.79 | 1688.16 | 1224.09 |
| 471 | 714.556 | 1345.51 | 1705.8 | 1235.93 |
| 472 | 718.158 | 1357.74 | 1711.76 | 1250.82 |
| 473 | 723.304 | 1367.25 | 1717.77 | 1264.65 |
| 474 | 717.138 | 1358.81 | 1716.96 | 1268.45 |
| 475 | 707.536 | 1357.76 | 1711.33 | 1267.09 |
| 476 | 708.993 | 1364.06 | 1711.81 | 1254.68 |
| 477 | 714.656 | 1361.43 | 1711.38 | 1253.9 |
| 478 | 711.792 | 1367.12 | 1714.54 | 1252.02 |
| 479 | 713.133 | 1367.06 | 1706.4 | 1257.68 |
| 480 | 705.717 | 1364.9 | 1693.19 | 1259 |
| 481 | 705.566 | 1368.24 | 1683.15 | 1248.85 |
| 482 | 698.52 | 1349.96 | 1675.53 | 1245.02 |
| 483 | 702.858 | 1346.45 | 1670.05 | 1239.84 |
| 484 | 704.507 | 1342.81 | 1678.88 | 1239.75 |
| 485 | 696.587 | 1340.95 | 1669.29 | 1233.55 |
| 486 | 698.11 | 1343.46 | 1664.56 | 1224.38 |
| 487 | 687.151 | 1328.42 | 1635.15 | 1216.81 |
| 488 | 688.581 | 1314.86 | 1630.7 | 1207.64 |
| 489 | 675.505 | 1307.44 | 1618.51 | 1191.37 |
| 490 | 664.987 | 1286.85 | 1604.83 | 1176.1 |
| 491 | 664.861 | 1278.77 | 1588.91 | 1153.81 |
| 492 | 670.804 | 1261.61 | 1579.75 | 1144.08 |
| 493 | 661.745 | 1247.61 | 1563.77 | 1137.18 |
| 494 | 645.43 | 1221.95 | 1533.58 | 1119.43 |
| 495 | 632.554 | 1206 | 1514.45 | 1102.37 |
| 496 | 624.36 | 1188.41 | 1499.42 | 1080.44 |
| 497 | 613.034 | 1160.86 | 1480.12 | 1062.26 |
| 498 | 602.661 | 1150.47 | 1456.73 | 1054.32 |
| 499 | 593.925 | 1136.7 | 1442.84 | 1037.87 |
| 500 | 588.507 | 1127.84 | 1415.37 | 1016.33 |
| 501 | 576.86 | 1107.34 | 1395.51 | 1004.19 |
| 502 | 565.906 | 1082.84 | 1377.47 | 988.742 |
| 503 | 547.317 | 1052.06 | 1348.94 | 970.458 |
| 504 | 537.43 | 1033.66 | 1323.14 | 950.514 |
| 505 | 528.484 | 1016.01 | 1290.86 | 928.377 |
| 506 | 519.338 | 988.333 | 1264.83 | 911.946 |
| 507 | 508.843 | 966.636 | 1231.9 | 889.221 |
| 508 | 496.326 | 943.318 | 1195.45 | 871.873 |
| 509 | 486.241 | 933.7 | 1170.98 | 853.053 |
| 510 | 471.114 | 907.892 | 1141.71 | 829.03 |
| 511 | 468.509 | 894.454 | 1117.38 | 809.058 |
| 512 | 473.699 | 877.338 | 1096.7 | 795.924 |
| 513 | 459.58 | 855.896 | 1063.02 | 776.138 |
| 514 | 458.353 | 833.097 | 1041.13 | 760.633 |
| 515 | 441.955 | 801.413 | 1014.58 | 736.768 |
| 516 | 429.32 | 778.943 | 980.18 | 714.754 |
| 517 | 415.438 | 745.346 | 944.189 | 694.883 |
| 518 | 396.472 | 718.392 | 910.299 | 669.975 |
| 519 | 392.267 | 700.691 | 888.96 | 647.859 |
| 520 | 387.272 | 684.353 | 866.074 | 627.735 |
| 521 | 380.199 | 674.866 | 848.37 | 616.733 |
| 522 | 373.952 | 659.82 | 838.301 | 604.27 |
| 523 | 369.678 | 650.618 | 814.836 | 594.78 |
| 524 | 364.948 | 638.228 | 801.501 | 579.77 |
| 525 | 356.604 | 616.843 | 780.812 | 563.789 |
| 526 | 354.293 | 597.741 | 760.044 | 554.361 |
| 527 | 344.546 | 586.766 | 748.54 | 543.4 |
| 528 | 328.768 | 566.423 | 722.004 | 520.859 |
| 529 | 320.666 | 548.377 | 703.132 | 508.697 |
| 530 | 310.201 | 538.233 | 684.886 | 493.716 |
| 531 | 303.975 | 533.559 | 668.735 | 487.386 |
| 532 | 297.389 | 524.305 | 658.482 | 476.217 |
| 533 | 290.109 | 510.276 | 642.413 | 463.518 |
| 534 | 276.295 | 499.752 | 625.993 | 443.382 |
| 535 | 284.62 | 491.67 | 612.184 | 437.639 |
| 536 | 276.288 | 478.937 | 599.848 | 427.726 |
| 537 | 260.508 | 457.771 | 590.678 | 416.847 |
| 538 | 252.759 | 439.372 | 566.955 | 403.242 |
| 539 | 239.839 | 419.964 | 548.916 | 384.28 |
| 540 | 223.467 | 395.854 | 521.976 | 365.57 |
| 541 | 209.482 | 387.382 | 503.008 | 357.462 |
| 542 | 192.465 | 366.514 | 476.832 | 341.488 |
| 543 | 184.9 | 348.519 | 453.094 | 321.42 |
| 544 | 189.305 | 341.968 | 436.846 | 308.442 |
| 545 | 175.087 | 323.212 | 419.289 | 301.362 |
| 546 | 163.627 | 311.207 | 399.983 | 291.868 |
| 547 | 155.943 | 288.914 | 381.896 | 281.633 |
| 548 | 154.073 | 265.784 | 367.082 | 266.1 |
| 549 | 146.544 | 255.125 | 341.6 | 247.996 |
| 550 | 139.49 | 243.916 | 334.346 | 234.992 |
| 551 | 129.593 | 227.482 | 314.173 | 226.456 |
| 552 | 117.676 | 218.968 | 299.974 | 215.692 |
| 553 | 120.245 | 213.453 | 286.64 | 208.577 |
| 554 | 121.931 | 211.437 | 272.626 | 206.554 |
| 555 | 117.532 | 202.888 | 263.274 | 193.378 |
| 556 | 112.399 | 198.842 | 253.095 | 190.18 |
| 557 | 116.48 | 195.911 | 246.536 | 186.603 |
| 558 | 112.702 | 184.95 | 232.852 | 181.876 |
| 559 | 104.351 | 177.262 | 218.003 | 176.277 |
| 560 | 108.677 | 171.584 | 219.7 | 177.077 |
| 561 | 105.849 | 164.911 | 221.252 | 180.153 |
| 562 | 107.814 | 164.19 | 221.49 | 176.179 |
| 563 | 105.436 | 158.726 | 218.69 | 167.665 |
| 564 | 98.6064 | 157.932 | 204.666 | 154.469 |
| 565 | 92.187 | 153.067 | 194.946 | 152.23 |
| 566 | 94.4773 | 150.51 | 192.12 | 154.704 |
| 567 | 95.8089 | 144.1 | 186.698 | 159.406 |
| 568 | 103.161 | 134.055 | 182.985 | 156.994 |
| 569 | 92.1617 | 122.386 | 176.277 | 150.125 |
| 570 | 79.9085 | 118.987 | 167.876 | 140.961 |
| 571 | 73.4409 | 109.32 | 153.306 | 138.509 |
| 572 | 67.9444 | 105.741 | 141.712 | 132.979 |
| 573 | 71.9476 | 97.5764 | 140.999 | 124.955 |
| 574 | 82.1635 | 105.113 | 149.798 | 125.489 |
| 575 | 79.6408 | 103.359 | 154.802 | 125.345 |
| 576 | 71.4367 | 97.6811 | 152.923 | 122.054 |
| 577 | 61.0527 | 88.768 | 134.214 | 120.913 |
| 578 | 55.9614 | 78.2746 | 117.003 | 107.195 |
| 579 | 48.8469 | 65.9492 | 104.803 | 96.7773 |
| 580 | 52.1117 | 67.4797 | 105.611 | 92.3075 |
| 581 | 50.7228 | 61.0254 | 106.152 | 90.2663 |
| 582 | 47.0493 | 58.9325 | 102.098 | 85.5515 |
| 583 | 46.361 | 58.2121 | 99.98 | 84.4447 |
| 584 | 40.9544 | 67.5376 | 100.965 | 77.9844 |
| 585 | 46.7078 | 70.8274 | 101.212 | 79.0225 |
| 586 | 51.1413 | 66.688 | 87.4162 | 71.6867 |
| 587 | 49.8742 | 62.6692 | 82.9145 | 69.6508 |
| 588 | 52.2914 | 57.6797 | 85.3184 | 60.5642 |
| 589 | 52.1519 | 49.5486 | 86.4113 | 63.1306 |
| 590 | 45.255 | 53.506 | 79.8761 | 63.1465 |
| 591 | 43.312 | 52.7457 | 70.1165 | 59.1625 |
| 592 | 35.4878 | 41.4448 | 57.2837 | 51.9939 |
| 593 | 39.2885 | 40.5558 | 59.5389 | 46.9044 |
| 594 | 44.4488 | 49.0506 | 69.4329 | 47.9623 |
| 595 | 44.4527 | 54.3647 | 67.8788 | 47.255 |
| 596 | 47.6226 | 56.5208 | 67.5738 | 54.1821 |
| 597 | 42.6912 | 50.9825 | 62.8766 | 55.6145 |
| 598 | 43.4553 | 49.1146 | 60.3639 | 53.359 |
| 599 | 41.1217 | 46.9166 | 53.8214 | 52.1713 |

**Fig S1:** pH dependence of bioluminescence activity: the measurement conditions are 2 μM of CTZ, 0.2 μM of GLuc, 0.2 M NaBr plus 50 mM MES (pH 6.0 or pH 7.0) or 50 mM MOPS (pH 7.9) or 50 mM Tricin (pH 8.8). The pH was adjusted with NaOH.

Light Intensity at 473 nm

| Time (s) | - VC |
| --- | --- |
| 0 | 519.936 |
| 8 | 499.75 |
| 20 | 531.272 |
| 41 | 478.236 |
| 86 | 286.954 |
| 121 | 209.279 |
| 156 | 149.093 |
| 180 | 132.648 |

| Time (s) | + VC |
| --- | --- |
| 0 | 477.464 |
| 14 | 561.811 |
| 32 | 652.147 |
| 61 | 562.19 |
| 102 | 402.221 |
| 134 | 303.909 |
| 162 | 249.601 |
| 194 | 210.912 |
| 242 | 172.081 |
| 292 | 139.006 |
| 322 | 106.738 |

| Time (s) | no salt |
| --- | --- |
| 0 | 85.2134 |

**Fig S2:**Time course of the luminescence after addition of Vitamin C: In the activity measurement, 1 μM of CTZ was luminesced in the presence of GLuc (0.25 μM) in 50 mM MES buffer containing 0.3M Ascorbic acid and 0.2 M NaCl, pH 7.0
